# Human Motor Cortex Encodes Complex Handwriting Through a Sequence of Primitive Neural States

**DOI:** 10.1101/2024.02.05.578548

**Authors:** Yu Qi, Xinyun Zhu, Xinzhu Xiong, Xiaomeng Yang, Nai Ding, Hemmings Wu, Kedi Xu, Junming Zhu, Jianmin Zhang, Yueming Wang

## Abstract

How the human motor cortex (MC) orchestrates sophisticated fine movements such as handwriting remains a puzzle^1–3^. Here, we investigate this question through Utah array recordings from human MC hand knob, during imagined handwriting of Chinese characters (306 characters tested, 6.3 ± 2.0 strokes per character). We find MC programs the writing of complicated characters by sequencing a small set of primitive states: The directional tuning of motor neurons remains stable within each primitive state but strongly varies across states. Furthermore, the occurrence of a primitive state is encoded by a separate set of neurons not directly involved in movement control. By automatically identifying the primitive states and corresponding neuronal tuning properties, we can reconstruct a recognizable writing trajectory for each character (84% improvement in reconstruction accuracy compared with baseline). Our findings unveil that skilled, sophisticated movements are decomposed into a sequence of primitive movements that are programmed through state-specific neural configurations, and this hierarchical control mechanism sheds new light on the design of high-performance brain-computer interfaces.

## Introduction

Humans are superb at controlling sophisticated fine movements, such as writing, typing, and musical performance^1,2^. These sophisticated motoric behaviors are often decomposed into a sequence of simpler movements^3–6^. For example, a word is decomposed into a sequence of letters, a letter can be further decomposed into strokes, and even a stroke may involve a complex movement trajectory. Such decomposition can reduce the complexity and shorten the time scale of each movement unit, which is of particular importance for the motor cortex (MC), since neurons in the MC generally show relatively simple tuning to movement features^7–12^, and the tuning varies over relatively long time scales^13^. It remains unclear, however, what are the primitive units during the programming of sophisticated fine movements and how each primitive unit is encoded in the MC.

Handwriting, a skill developed through years of deliberate practice, serves as an example of sophisticated fine motor control in humans. This study investigated the neural basis of handwriting by recording single-unit neural activity from human MC with two 96-channel Utah microelectrode arrays in the left hand knob area of the precentral gyrus^14^ (Fig. 1a and Extended Data Fig. 1a). We studied imagined writing of Chinese characters (Fig. 1b, c), which are highly complex (> 3500 frequent characters composed by 32 types of strokes) and pose a great challenge for motor control. We identified that, during the writing of a character, the directional tuning of neurons alternates among a few primitive states, each controlling the writing of a fragment of a character (Fig. 1d-g). Furthermore, the occurrence of each primitive state is signified by a set of state-occurrence neurons that do not show directional tuning (Fig. 1g). Computational models that decompose the writing process into a sequence of primitive states better explain spiking activity from individual neurons (279% change) and better decode the writing trajectory based on the neural population (84% change, Fig. 1h, i), compared to a model that assumes stable neuronal tuning throughout the writing of a Chinese character.

**Fig. 1.**
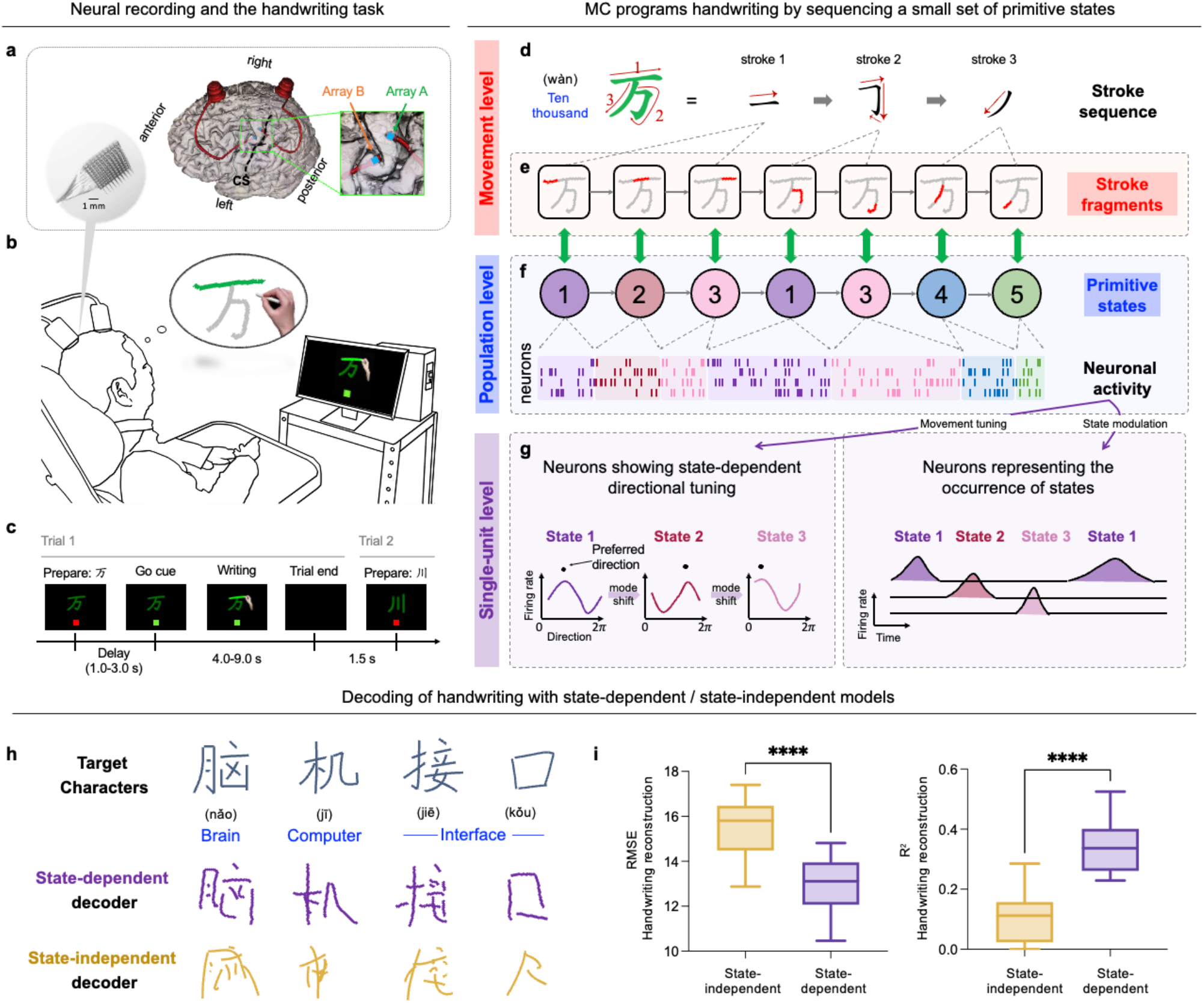
The handwriting task and neural representation during imagined handwriting. **a**, The participant has two 96-channel Utah intracortical microelectrode arrays implanted in the left motor cortex. Both arrays (array-A and array-B) are positioned in the hand knob area, approximately 2 mm apart from each other. **b**, The participant is instructed to imagine handwriting under video guidance (Extended Data Video 1). **c**, Trial design schematic. Each trial consists of a single character displayed on the screen. A trial begins with the target character being displayed on the screen (prepare), followed by a go cue (green light), and acted writing of the character stroke by stroke (see Methods). **d**, Writing of an example Chinese character in terms of strokes (Top). **e-g**, Summary of the main findings. MC programs character the writing by sequencing a small set of primitive states (**f**), each encoding a specific stroke fragment (**e**). A set of neurons exhibits directional tuning that is stable within each primitive state but strongly variable across states (**g**, left). Another set of neurons is not directly involved in movement control but represents the occurrence of states (**g**, right). **h**, The writing trajectory was decoded using state-dependent and state-independent neural decoders. **i**, Performance of handwriting decoding using Root Mean Square Error (RMSE) and R^2^ between the ground truth trajectory and the neurally decoded trajectory (P < 10^-4^, Wilcoxon matched-pairs signed rank test, n = 17 sessions).

### Evidence of state-dependent encoding during handwriting

The participant imagined handwriting Chinese characters under the guidance of a video (Fig. 1a-c, Extended Data Fig. 1a-b, and Extended Data Video 1). We recorded 2850 neurons across 20 experimental sessions (see Supplementary Information), and each character was written 3 times. We first analyzed whether some neurons reliably responded during the writing of individual characters using the Fisher’s discriminant value (FD), which was higher if a neuron generated similar responses when writing the same character twice than when writing two different characters (see Supplementary Information, Fig. 2a). Hundreds of neurons showed FD much higher than the chance level, i.e., 1.0 ± 0.05 (M ± SD, estimated by shuffling the responses across characters), demonstrating reliable responses during character writing. We next analyzed whether the reliable neural responses to characters could be explained by the classic velocity-based directional-tuning model^8,9,15^. The reliability of neural response to individual characters, quantified by the FD, only had a moderate correlation with neural sensitivity to velocity direction, quantified by the *R*^2^ of the directional-tuning model (*R* = 0.46). In other words, neuronal tuning to velocity direction could not well explain neural responses to characters. In the following, to facilitate discussion, we loosely defined two categories of neurons: Neurons not well explained by the directional-tuning model were referred to as the complex-tuning neurons (n = 115 under the criterion that *R*^2^ < 0.1 and FD ≥ 1.4, shown in blue in Fig. 2a), and neurons well explained by the directional-tuning model were referred to as simple-tuning neurons (n = 64 under the criterion that *R*^2^ ≥ 0.1 and FD ≥ 1.4, shown in magenta in Fig. 2a).

**Fig. 2.**
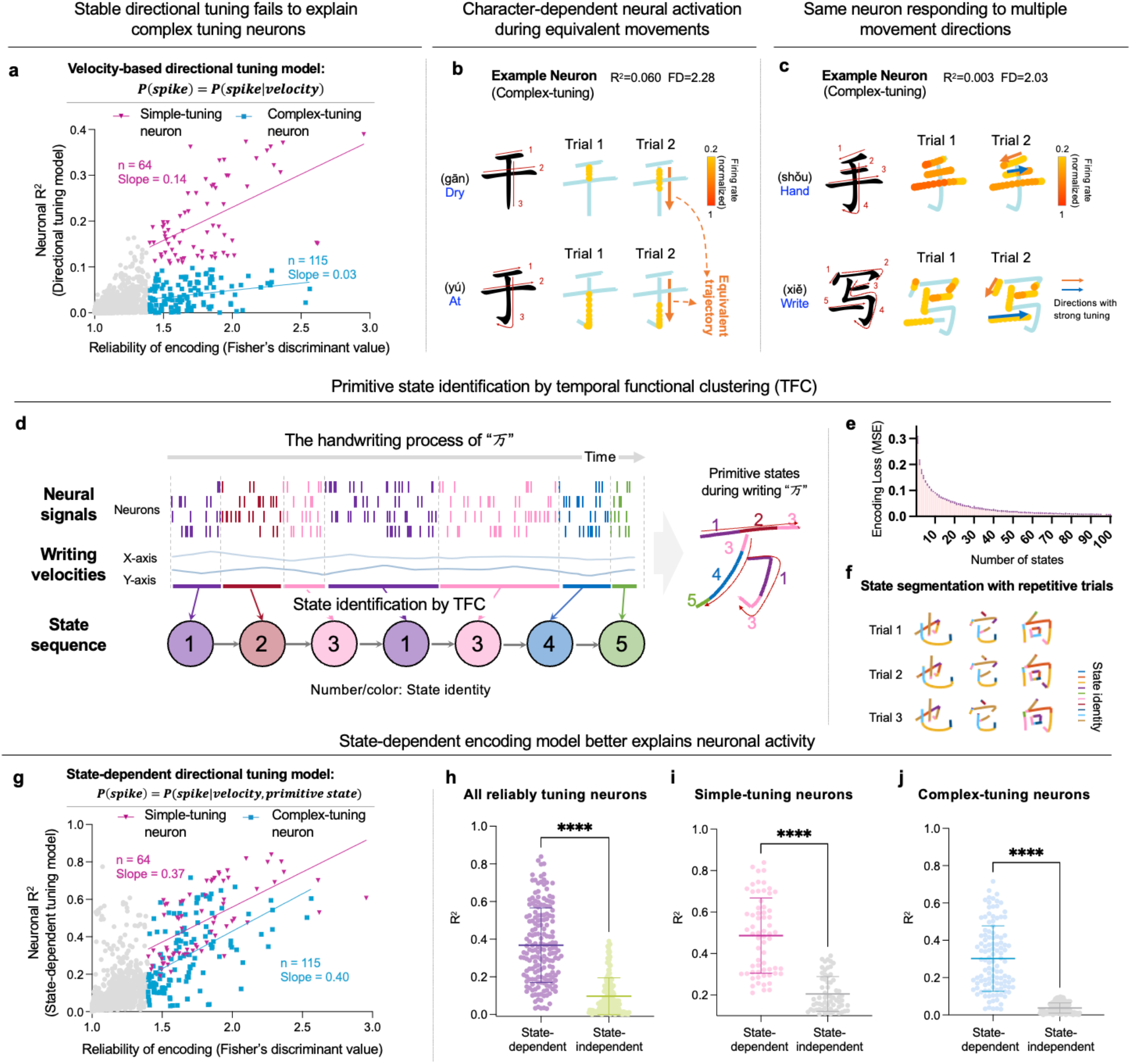
State-dependent directional tuning model better explains neural activity during handwriting. **a**, Reliability and tuning property of neurons (N = 2850 neurons across 20 sessions). The reliability of the response to each character is characterized using the Fisher’s discriminant value (FD) across characters (the 30 within each session), while the tuning property is characterized using the R^2^ of the directional tuning model. Neurons with reliable tuning but cannot be explained by the directional tuning model are referred to as complex tuning neurons (n = 115, R^2^ < 0.1 and FD≥1.4), and neurons conformed to the directional tuning model are referred to as simple tuning neurons (n = 64, R^2^≥0.1 and FD≥1.4). **b-c**, Responses of exemplary complex-tuning neurons during handwriting. The firing rate in each 50 ms time bin is visualized by the color of a dot, on the trajectory of the character being written. In (**b**), the neuron shows diverged responses to the same downward movement in two characters. In (**c**), the neuron can respond to multiple movement directions but does not consistently respond to any direction. **d**, Neural activity recorded during imagined handwriting is divided into nonoverlapping primitive states, which are classified using the temporal functional clustering (TFC) algorithm. Each color denotes a specific primitive state. **e**. The model encoding loss decreases when more primitive states are modeled, and the tipping point occurs around 10 states. **f**. Repetitive handwriting trials were divided into consistent sequences of primitive states. **g-j**. R^2^ of the model that explains neural activity using state-dependent directional tuning. The state-dependent directional tuning model significantly enhances the R^2^ for neurons with reliably tuning to characters (P < 10^-4^, Wilcoxon matched-pairs signed rank test, n = 179), and the mean R^2^ increased from 0.097 to 0.368 (**h**). For simple-tuning neurons, the mean R^2^ increased from 0.202 to 0.487 (**i**, P < 10^-4^, Wilcoxon matched-pairs signed rank test, n = 64). For complex-tuning neurons, the mean R^2^ increased from 0.038 to 0.302 (**j**, P < 10^-4^, Wilcoxon matched-pairs signed rank test, n = 115).

To investigate why complex-tuning neurons were not well explained by the directional-tuning model, we first illustrated the responses of complex-tuning neurons by overlaying neural firing rate on the trajectory of characters and observed two general phenomena. First, some neurons reliably responded during the writing of a small fragment of a character, but did not respond during the writing of other fragments that involved movements in the same direction. For example, the same downward vertical movement was involved when writing two characters, i.e., ‘干’ and ‘于’, which have highly similar orthographical forms but different pronunciations and meanings. A complex-tuning neuron, however, responded differently during the same continuous downward movement in the two characters: It responded during the beginning of the movement for one character but responded near the end of the movement for the other character (Fig. 2b). Second, some neurons responded to different movement directions in different character fragments. For example, the same neuron may respond to leftward (e.g., stroke 1 in ‘手’), rightward (e.g., stroke 2 in ‘手’), and downward (e.g., first portion of stroke 3 in ‘写’) movements when writing some fragments of a character (Fig. 2c), but do not respond to the same movement directions when writing other fragments (e.g., first portion of stroke 2 in ‘写’ for rightward movement, first portion of stroke 4 in ‘手’ for downward movement).

### Primitive states and state-dependent tuning

Previous analyses have shown that the directional tuning of the neuron might change when writing different fragments. Based on these findings, we hypothesized that the writing of a character was decomposed into a sequence of primitive states: Only within each primitive state, movement was controlled according to the classic directional tuning model, while different primitive states employed different directional tuning models. This model was formulated as

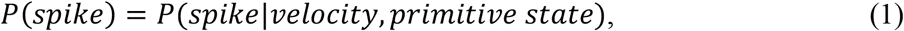

where *P*(*spike*) is the probabilistic distribution of the firing rate. To identify the primitive states, we developed an algorithm, termed Temporal Functional Clustering (TFC; Fig. 2d, Supplementary Materials), which grouped the movement tuning in each time bin (50 ms in duration) into a few clusters under a temporal continuity constraint (see Extended Data Fig. 2 for verification of the algorithm).

A hyperparameter of the model was how many primitive states were allowed. When only one state was allowed, the model reduced to the classic directional tuning model, and the model complexity increased when the number of states increased. We found that a small number of primitive states was enough to significantly better predict the neural responses (P < 10^-3^ for each state number ≥ 2, Wilcoxon matched-pairs signed rank test, Fig. 2e) and, as a tradeoff between model performance and model complexity, we chose to model the character writing process with 10 primitive states in the current study. Note that the number of states used here was much smaller than the number of Chinese characters (n = 306) or even the types of strokes for Chinese characters (n = 32).

The state-dependent model decomposed the writing of a character into a sequence of primitive states, and we observed that a character was decomposed into a similar state sequence in multiple trials (Fig. 2f), indicating that the primitive state sequence was stored in long-term memory, instead of being composed online during movement. We found that the state-dependent model better explained the neural responses of individual neurons than the classic directional-tuning model (P < 10^-4^, Wilcoxon matched-pairs signed rank test, Fig. 2g, h), for both simple-tuning neurons (Fig. 2i) and complex-tuning neurons (Fig. 2j). For complex-tuning neurons, the mean *R*^2^ increased from 0.04 to 0.30 (P < 10^-4^, Wilcoxon matched-pairs signed rank test). In the following, we explored why the state-dependent model improved the modeling of simple- and complex-tuning neurons.

First, we observed that the directional tuning models learned from one primitive state could well explain neural activity during that primitive state but not other primitive states (Fig. 3a). For each neuron, the state-dependent model estimated a directional tuning curve per primitive state, and we illustrated these tuning curves for both simple- and complex-tuning models (Fig. 3b for normalized tuning curves of all neurons, Fig. 3c-d for tuning curves of representative neurons, and Extended Data Fig. 3 for more examples). For simple-tuning but not complex-tuning neurons, the tuning curves had similar shapes across different primitive states (Fig. 3b, c). To quantify this effect, we analyzed the preferred direction of each tuning curve (PD, representing the direction with the largest firing rates) and the standard deviation of PD was higher for complex-tuning than simple-tuning neurons (P < 10^-4^, Mann-Whitney test, Fig. 3e). Next, we further analyzed the modulation depth of each tuning curve (MD, defined as the peak-to-peak value of the directional tuning curve). It was found that the MD varied across primitive states even for simple neurons and the standard deviation of MD was comparable for complex- and simple-tuning neurons (P = 0.08, Mann-Whitney test, Fig. 3f). The state-dependent MD could explain why the state-dependent model outperformed the state-independent model even for simple-tuning neurons.

**Fig. 3.**
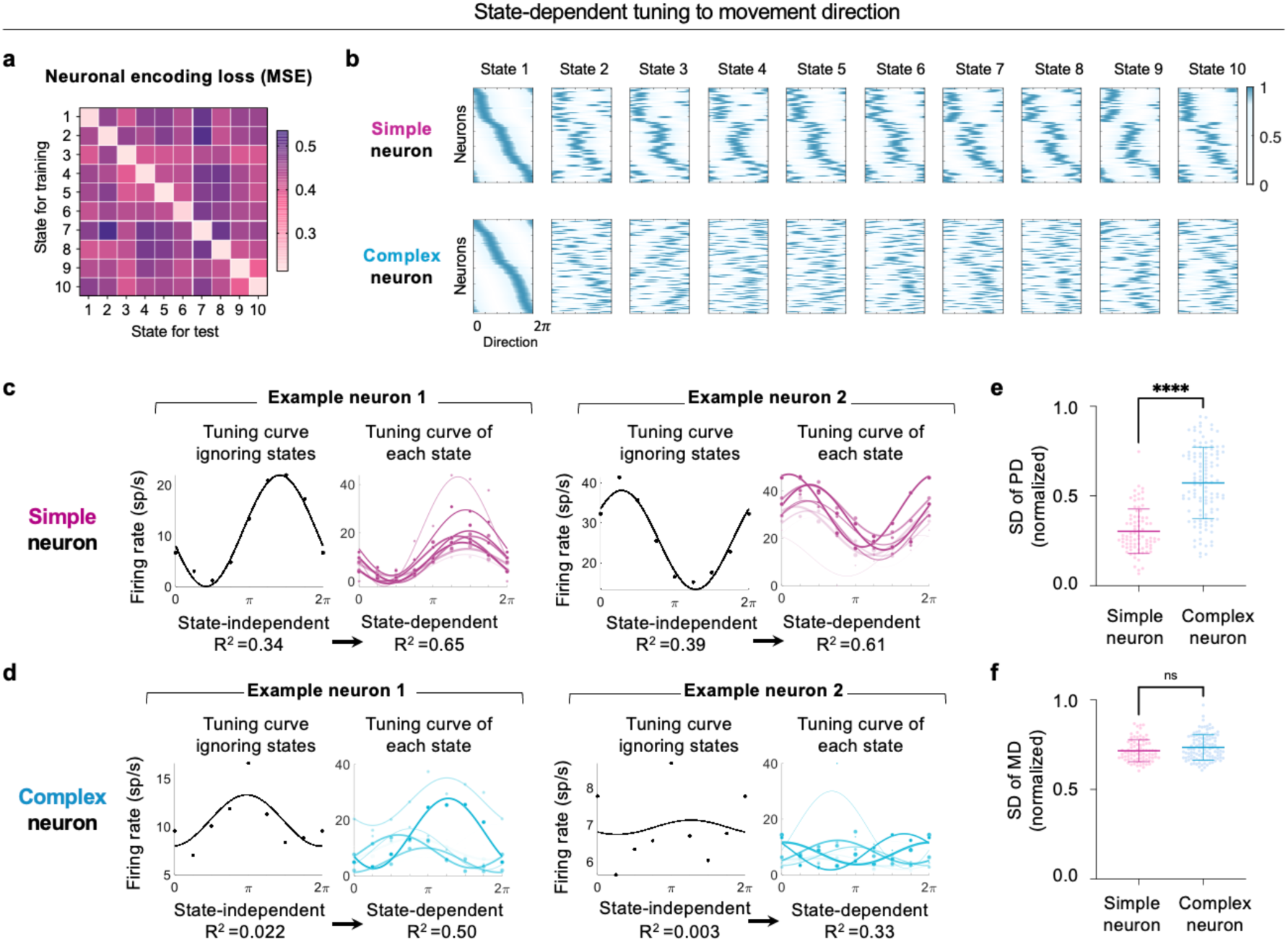
State-dependent directional tuning of MC neurons. **a**, Pairwise neural activity encoding loss between primitive states. Each matrix entry (*i, j*) indicates the prediction error of neural activity in primitive state *j* for the model trained on primitive state *i*. A model can only well predict the neural responses during the primitive state it is trained on. **b**, Normalized firing rates over directions with both simple- and complex-tuning neurons, sorted based on state 1. The Simple tuning neurons show more consistent directional tuning over primitive states compared with the complex-tuning neurons. **c**, Directional tuning curve of two exemplar simple tuning neurons estimated using a state-independent model (black) or a state-dependent model (magenta). For the state-dependent model, a tuning curve is estimated based on each primitive state and the tuning curves have a consistent preferred direction (PD) across primitive states but the modulation depth (MD) varies across primitive states. Darker color for curves with higher R^2^. **d**, Similar to (**c**), but for complex-tuning neurons that exhibit variable directional tuning under different primitive states. **e-f**, Statistical analysis of PD (**e**) and MD (**f**) for simple- and complex-tuning neurons. The standard deviation of PD is significantly higher for complex-tuning neurons (mean = 0.57) than simple-tuning neurons (mean = 0.30) (P < 10^-4^, Mann-Whitney test).

### Neurons signifying the occurrence of primitive states

We demonstrated that the tuning properties of neurons varied across primitive states but it remained unclear how these primitive states per se were encoded. We hypothesized that a group of neurons in the MC controlled or signaled the occurrence of each primitive state and this group of neurons were not directly involved in movement control. To test this hypothesis, we analyzed neurons that showed reliable responses to characters, since the same character was divided into reliable state sequences across trials, but could not be explained by the state-dependent tuning curves (Fig. 4a, shown in purple). To quantify whether these neurons were not sensitive to movement features at all or whether they encoded features in a highly nonlinear way, we estimated the mutual information between neural activity and movement features including position, velocity, and acceleration. The mutual information was not much higher than chance (Fig. 4b), confirming that these neurons did not directly encode movement features. To test whether these neurons were tuned to specific primitive states, we calculated the mean firing rate of these neurons within each primitive state and found that these neurons fired during a small number of states (Fig. 4c). Therefore, these neurons were referred to as state-modulation neurons. We further illustrated the time course of the neuronal firing of state-occurrence neurons in Fig. 4d. It was observed that the response of state-occurrence neurons generally followed a stereotypical bell-shaped form, during the occurrence of specific primitive states.

**Fig. 4.**
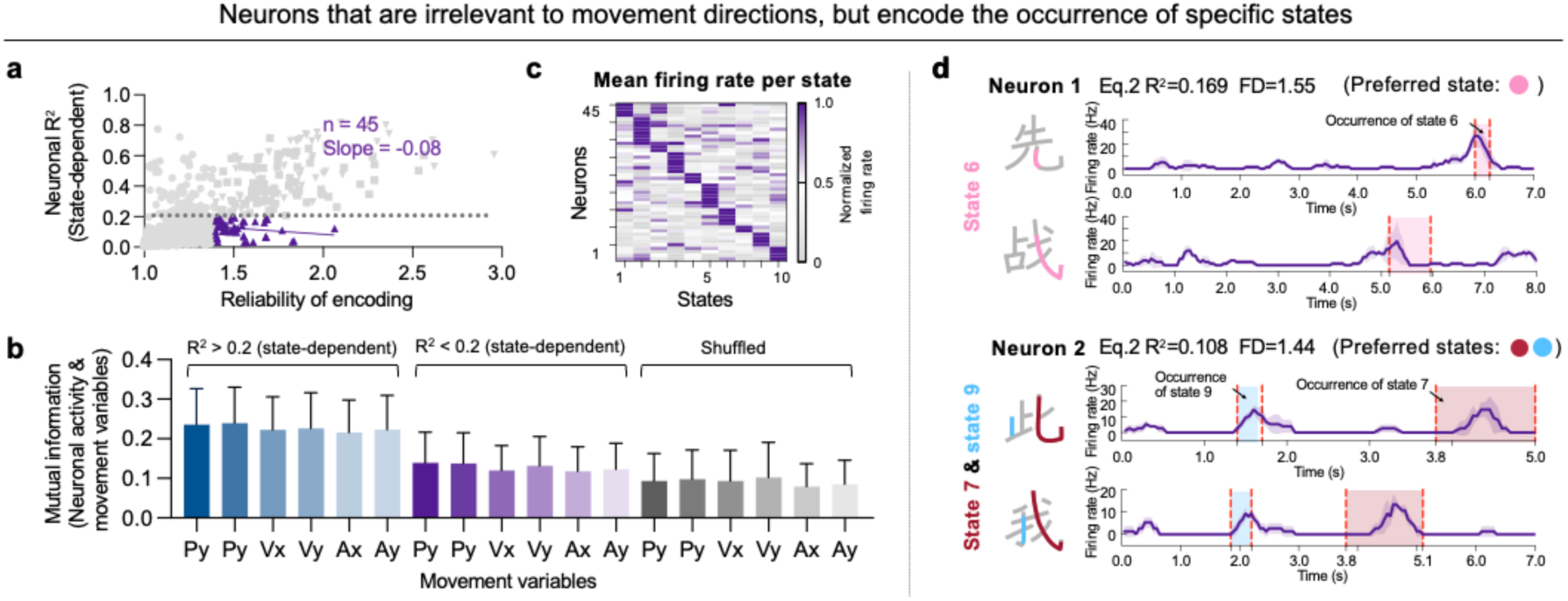
Neurons that indicate the occurrence of primitive states. **a**, A subset of complex-tuning neurons is not well explained even with the state-dependent tuning model (R^2^ < 0.2, shown in purple). **b**, Mutual information between neural activity and movement variables (position/P, velocity/V, and acceleration/A, over *x* and *y* axes). Neurons with R^2^ below 0.2 showed comparable dependence on movement variables compared with the shuffled responses. **c**, Normalized firing rates of neurons (R^2^ < 0.2) show high selectivity to primitive states. Purple colors represent high firing rates, indicating neurons that were active above the mean. Gray colors denote low firing rates below the mean. **d**, Response of exemplar neurons, which generates a bell-shaped response to the occurrence of a specific primitive state. The temporal firing rate during the handwriting process is averaged with three repetitions, with the shaded regions showing the standard deviation. The first exemplar neuron specifically responds to state 6 (shown in pink), and the second exemplar neuron tunes to both state 7 and state 9.

### Handwriting decoding based on primitive states

Previous analyses demonstrated how individual neurons encode the handwriting process, and in the following we utilized the population response to decode the handwriting trajectory (Fig. 1h). We built a state-dependent neural decoder to decode the movement velocity and recovered the handwriting trajectory based on velocity^14^ (Extended Data Fig. 4a). Compared with a state-independent neural decoder, the state-dependent decoder could decode handwriting with lower Root Mean Squared Error (RMSE, 10%-27% decrease, P < 10^-4^, Wilcoxon matched-pairs signed rank test, Fig. 1i) and higher *R*^2^ (> 84% increase, P < 10^-4^, Wilcoxon matched-pairs signed rank test, Fig. 1i). We further used a commercially available Optical Character Recognition (OCR) system^16^ to test the readability of decoded trajectories (Extended Data Fig. 4b), which output the top 10 candidate characters for each decoded trajectory. Within a pool of 400, 3500, and 96000 characters, the Top-1 candidate was correct for 90.0%, 87.8%, and 72.2% of characters, respectively, for the state-dependent decoder (Extended Data Fig. 4c). The decoding process could be achieved online, allowing the participant to use a robotic arm to write recognizable Chinese characters through imagination (Extended Data Video 2). Taken together, these results demonstrated that complex handwriting is decomposed into smaller units that are encoded in different primitive states in the MC, and only within each primitive state is movement encoded in a roughly linear manner.

## Discussion

Handwriting is a highly sophisticated skilled behavior that is special to the human being, and the inherent complexity of the Chinese writing task provides a unique window to dissect the underlying neural mechanisms for sophisticated fine movements. Through the task, we found that the human MC decomposes a sophisticated writing process into a sequence of primitive states, under the control of three types of neurons: (1) simple-tuning neurons that have stable directional tuning, (2) complex-tuning neurons that have variable directional tuning across primitive states, and (3) and neurons that encode the occurrence of each primitive state.

These three types of neurons can support hierarchical control of sophisticated movements (Fig. 5): A highly complex movement trajectory is first decomposed into small and simple fragments, and each trajectory fragment is further converted into a sequence of movement velocity under a specific primitive state of the MC. In this process, it is possible that (1) the state-occurrence neurons serve as “timestamps” for each primitive state, (2) the complex-tuning neurons are upstream motor neurons related to the encoding of trajectory fragments, and (3) the simple-tuning neurons are downstream motor neurons related to the control of movement velocity. This view is in sharp contrast with a view that the MC only carries out simple motor commands: The orthographic form of a character is decomposed into movement velocity required at each time moment outside the MC, and the MC only implements the motor command, i.e., the desired direction^7^ or velocity^17^, through simple-tuning neurons.

**Fig. 5.**
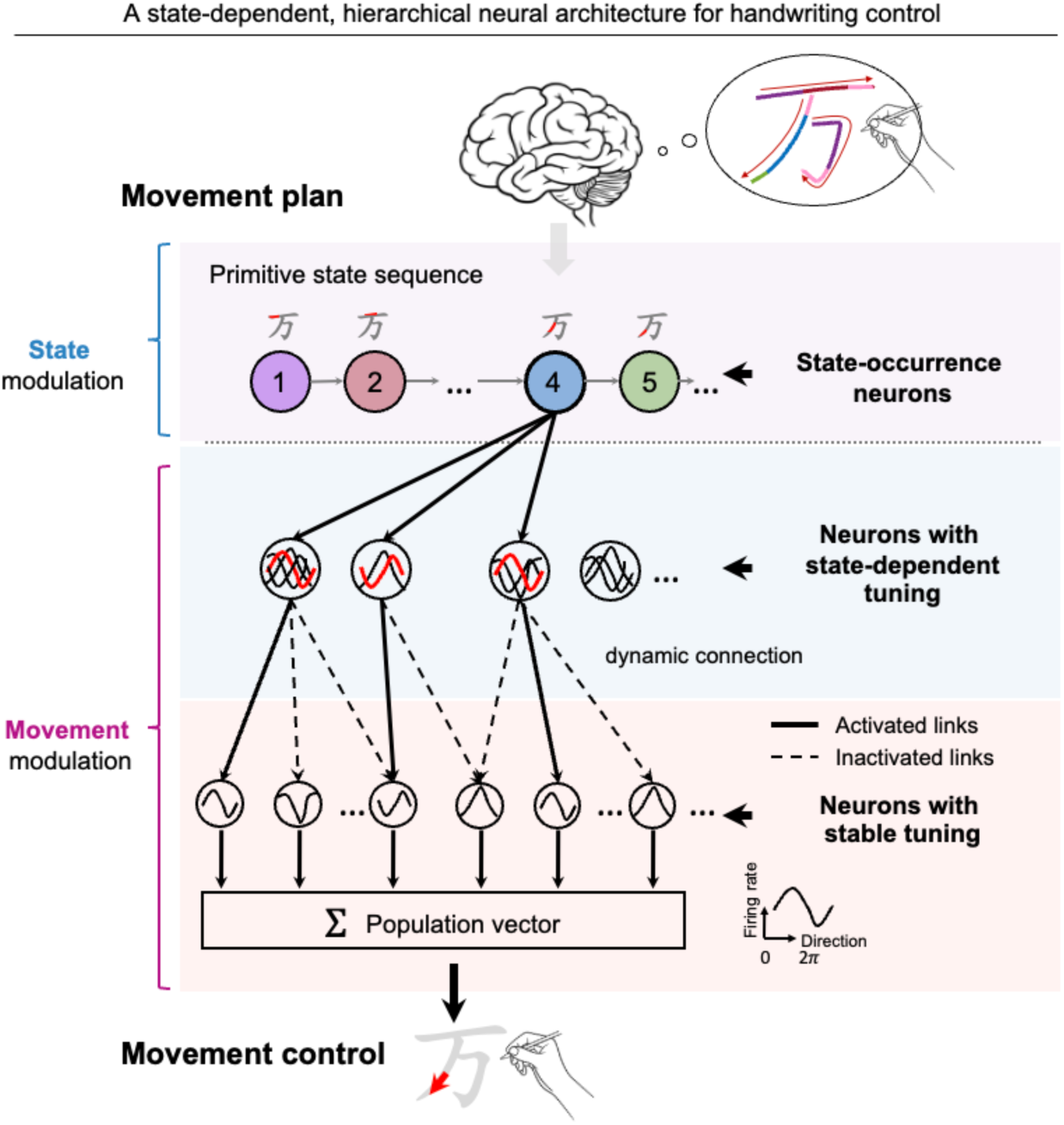
Neural processing architecture of handwriting. Sophisticated handwriting is divided into primitive states, each corresponding to a stroke fragment. Under each primitive state, movement is controlled through a state-specific neural ensemble, which possibly consists of three types of neurons. The state-occurrence neurons lead to or signal the occurrence of each primitive state. The complex tuning neurons encode a fragment of the movement trajectory and are connected to a set of simple tuning neurons, which directly control the movement direction at the current time moment.

Hierarchically decomposing a complex sequence into smaller units is a common strategy in the brain^5^ and its neural underpinnings have been studied across multiple domains such as motor planning^3^, decision making^18^, working memory^19^, song sequencing^20^, and language processing^21^. Revealing the primitive units to decompose a sequence, however, turns out to be highly challenging, especially for language-related processes^6,22^. Here, we proposed that the brain controls complex movements through a sequence of primitive states. Based on the writing behavior, characters and strokes are apparent candidates for the primitive unit for writing. Nevertheless, Chinese has a large number of characters (>3500 frequent characters), and assigning a specific neural population for the writing of each character, i.e., one-hot coding, is implausible. The number of stroke types is limited (n =32). Nevertheless, strokes can be sophisticated, such as ‘㇈’ and ‘㇋’, and redundant, i.e., sharing common elements such as vertical or horizontal movements. Furthermore, even the writing of a simple stroke such as ‘一’ can be meaningfully divided into a start, a body, and an end, which are especially emphasized in Chinese calligraphy and correspond to the decomposition of a movement into acceleration and deceleration phases^23^. Here, we demonstrate that, in the MC, a character is decomposed into units that correspond to fragments of a stroke, to possibly reduce the redundancy between strokes and to precisely control the start and end of a stroke.

Previous studies have demonstrated that neurons in the MC are tuned to movement features^7,8,11,24–26^ but static tuning to movement features has limited ability to explain the variance of neuronal activity^13,23,27^, especially during natural movements^28^. Instead, neural tuning to movement features can actively adapt to a specific task or environmental setting^29,30^. In other words, depending on the external environment or instruction, the MC neurons can encode movements through different neural encoding subspaces^31–33^. The current study, however, demonstrates intrinsic alternation between MC neural encoding subspace, depending on internally generated states during sophisticated movements. Similar ideas have also been proposed based on relatively simple movements, e.g., decomposing a reaching movement into an acceleration phase and a deceleration phase^13,23,34^ or stereotyped movement fragments^12^, but here we demonstrated many diverse primitive states, which can encode various movement fragments using under a linear directional-tuning model, and proposed a hierarchical model for the neural control of sophisticated movements (Fig. 5).

In summary, our results strongly demonstrate that sophisticated fine movements such as handwriting are encoded in the human MC by a sequence of primitive states, each controlling a fragment of the movement, and the state-dependent encoding mechanism can shed light on future design of BCIs for sophisticated fine movements.

## Methods

### Participant and ethics

All clinical and experimental procedures conducted in this study received approval from the Medical Ethics Committee of The Second Affiliated Hospital of Zhejiang University (Ethical review number 2019-158, approved on 05/22/2019) and were registered in the Chinese Clinical Trial Registry (Ref. ChiCTR2100050705). Informed consent was obtained verbally from the participant, along with the consent of his family members, and was duly signed by his legal representative.

The volunteer participant is a right-handed man, 75-year-old at the time of data collection. He was involved in a car accident and suffered from complete tetraplegia subsequent to a traumatic cervical spine injury at the C4 level, which occurred approximately two years prior to study enrollment. The volunteer participant demonstrated the ability to move body parts above the neck and exhibited normal linguistic competence and comprehension for all tasks. He scored 0/5 on skeletal muscle strength for limb motor behavior.

On August 27, 2019, two 96-channel intracortical microelectrode arrays (4 mm × 4 mm Utah Array with 1.5 mm length, Blackrock Microsystems, Salt Lake City, UT, USA) were implanted in the left primary motor cortex, with one array located in the middle of the hand-knob area (array-A) and the other array located medially approximately 2 mm apart (array-B), guided by structural (CT) and functional imaging (fMRI). The participant was asked to perform imagery movement of hand grasping and elbow flexion/extension with fMRI scanning to confirm the activation area of the motor cortex. Data presented in this study cover the period from post-implant days 1374 to 1792.

### Neural signal recording and processing

Neural signals were recorded from the microelectrode arrays using the Neuroport^TM^ system (NSP, Blackrock Microsystems). The signals were amplified, digitized, and recorded at a sampling rate of 30 kHz. To reduce common mode noise, a common average reference filter was applied, subtracting the average signal across the array from each electrode. A digital highpass filter with a cutoff frequency of 250 Hz was then applied to each electrode. Then, threshold crossing detection was performed using the Central software suite (Blackrock Microsystems). The threshold was set based on the root mean square (RMS) of the voltage time series recorded on each electrode. Specifically, thresholds ranging from −6.25×RSMto −5.5×RSMwere used.

To analyze the neuronal activity, neurons were manually sorted with either Plexon Offline Spike Sorter v4 (offline analysis) or Central software (online decoding). After spiker sorting, neuronal spikes were binned into 50 ms bins, without overlapping. For the decoding tasks, the spike data were smoothed using an average filter of 5 bins.

The delay between the start of data extraction and the go cue was determined by evaluating a set of delay values ranging from −1,000 ms to 1,000 ms. The delay value of 300 ms was selected according to the overall decoding performance with a linear model.

### Experimental paradigm

The participant performed an imagined handwriting task. During the experimental sessions, the participant imagined the handwriting of characters guided by instructional videos (Fig. 1 and Extended Data Video 1). The instructional video presented a virtual hand writing a target character stroke by stroke; at the same time, the participant imagined writing the character as if the virtual hand belonged to him.

For the Chinese character writing task, there were two paradigms.

1) Single-character writing with visual guidance: a single Chinese character in dark green color was displayed on the screen above a red square during the delay period, which lasted between 1 to 3 seconds. After the delay period, the red square cue turned green and played a sound ‘ding’ to signal the participant to start writing. Simultaneously, a virtual hand holding chalk appeared on the screen and wrote the character stroke by stroke, highlighting the written part in bright green. The duration of the writing period varied depending on the complexity of the Chinese character, with more strokes requiring a longer writing time. After completing one character, the screen turned black and lasted for 1.5 seconds (please see Extended Data Video 1).
2) Sentence writing with visual guidance: initially, the characters of a sentence were displayed in dark green on the screen above a red square. Following a delay period, the red square turned green, indicating the start of the sentence. The experimental procedure for each character within the sentence was identical to the visual guidance paradigm, with the exception that there was no ‘ding’ sound during the writing, but a short horizontal line appeared below the character as a reminder for the participant to begin writing (please see Extended Data Video 2).

The instructional video presents the writing process in a stroke-by-stroke manner. To facilitate smooth tracking, the velocity of the virtual hand was programmed to maintain a constant acceleration, ensuring ease of following for the participant.

The online neural signal processing program was developed in MATLAB. The graphical user interface (GUI) of the experimental task was developed in Python. Offline signal processing and analysis systems were developed by MATLAB and Python.

### Temporal functional clustering (TFC) algorithm

The temporal functional clustering (TFC) algorithm is a computational approach designed to identify state and state switches during a writing process. The underlying assumption of TFC is that the neuronal encoding mode, at the population level, may shift under different states while keeping stable within the same state. Under this assumption, TFC can automatically compute the neural mapping model for each state, and detect state switches in a data-driven way.

Specifically, given a paired dataset {*X*,*Y*}≡{*x*_1_,…, *x*_*T*_, *y*_1_,…, *y*_*T*_}, where 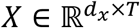 represents the writing kinematic data, and 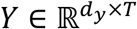 represents the preprocessed neural data, with *d*_*x*_ and *d*_*y*_ denoting the respective dimensions, and *T* representing the length of time bins.

Our objective is to learn *M* encoding models of *ℋ*_*m*_(·)∈ {*ℋ*_1_(·), *ℋ*_2_(·),…, *ℋ*_*M*_(·)}, where each model represents a single state.

To obtain the *M* encoding models, the TFC algorithm employs an Expectation-Maximum (EM) process. Initially, we randomly set the parameters of *M* encoding models. Then we perform the Expectation-step (E-step), where each pair of {*X*,*Y*} is assigned to a specific encoding model where the data has the lowest encoding loss. Especially, the encoding loss is smoothed temporally to encourage that the adjacent data pair should belong to the same state (temporal constraint). After that, we perform the Maximum-step (M-step), where each model updates its parameters with the data pairs assigned to it. The TFC algorithm repeats the E-step and M-step iteratively until convergence.

1) Initialization step To obtain *M* initial models, we randomly set the parameters of each model *ℋ*_*m*_(·). Each model is an encoding function that maps kinematics to neural signal estimation *Ŷ*:

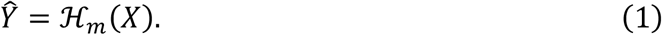
2) E-step: data assignment The E-step aims to reassign data pairs to their fittest models. For each data pair {*X*,*Y*}, we can compute the predicted neural signal *Ŷ*_*m*_ given kinematics *X* and encoding model *ℋ*_*m*_(·):

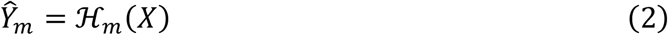

where 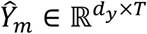 denotes the neural signals predicted by the encoding model *ℋ*_*m*_(·)for all the kinematics *X*. Given a set of *M* encoding models, we can obtain *M* predictions for each data pair. Then we calculate the encoding loss at each time step for each encoding model:

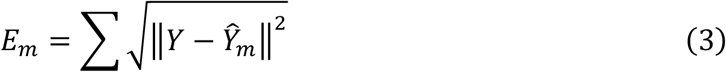

where *E*_*m*_ ∈ *ℝ*^1×T^ is the encoding error between the neural signal *Ŷ*_m_ estimated by model *ℋ*_*m*_(·), and the ground truth *Y*, summed over all the neurons. Next, we smooth the error vector *E*_*m*_ temporally to encourage data pairs that are adjacent in time to be assigned to the same model. With this constraint, we can obtain more temporally continuous state segmentation. The window size for a moving averaging smooth was typically set with 3 to 10 bins, denoted as *l*_smooth_. The smoothed encoding loss for model *ℋ*_*m*_(·)is given by:

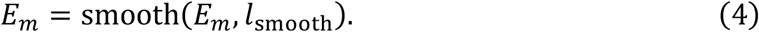 Once we have the smoothed encoding loss for each model, we assign each data pair to the model that gives the minimal encoding loss:

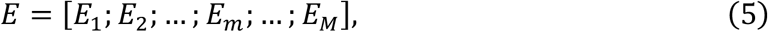

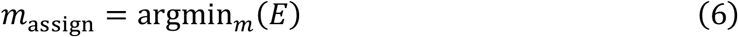

where *E* ∈ *ℝ*^*M*×*T*^ represents each model’s encoding loss, and *m*_assign_ ∈ *ℝ*^1×*T*^ denotes the model index selected.
3) M-step: parameter updating After the E-step, each data pair has been assigned to a specific model, and each model *ℋ*_*m*_(·)has a collection of data pairs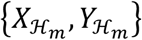. Then we can update the parameters for the models, by fitting the following function:

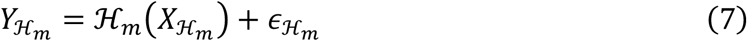

with 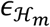 being the zero-Gaussian noise term. In this study, we use a linear function, such that *ℋ*_*m*_(·)would include a linear mapping matrix and a Gaussian noise.
4) Repeat E-step and M-step until convergence By iteratively performing the E-step and M-step, the overall encoding error will decrease continuously. Suppose at the *i*^*th*^ iteration we have an error of *e*_*i*_. One pre-set early-stopping threshold *β* (typically set to 0.001) can be used to decide when to stop the iteration:

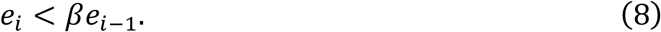

When the iteration stops, the TFC algorithm will return the parameters for each model (state) as well as the time indices corresponding to each model. The time indices indicate the segmentation of states.

### DyEnsemble neural decoder

Given a pool of encoding models learned by TFC, we can utilize the DyEnsemble decoder, which is a dynamic encoder-based decoder, to assemble state-specific encoding models during a writing process, adaptively to the change of states^2,3^.

The DyEnsemble decoder defines the state-space model as follows:

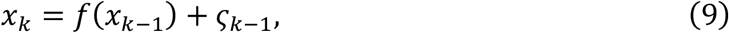

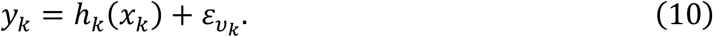

The model contains a state equation (Eq. 9) that transmits the state of interest *x*_*k*−1_ at time *k* − 1 to the next time step *k* using function *f*(·), with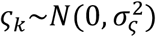), a zero-mean Gaussian, being the transition noise. It also contains an observation equation (Eq. 10), which uses *x*_*k*_, the state at time *k*, to infer the measurement variable *y*_*k*_ using function *h*_*k*_(·), with a zero-mean Gaussian observation noise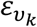. Notably, the observation function *h*_*k*_(·)is dependent on time, such that it dynamically changes over time. The DyEnsemble algorithm aims to estimate the dynamic measurement function (*h*_*k*_(·)), and the corresponding state (*x*_*k*_) over time, given the incoming neural signals (*y*_*k*_).

In the scenario of neural decoding, 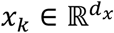 is the movement kinematics, and 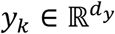 denotes the neural signals, with *d*_*x*_ and *d*_*y*_ denoting the respective dimensions. The observation function *h*_*k*_(·)is an encoding function mapping from kinematics (*x*_*k*_) to neural signals (*y*_*k*_). DyEnsemble model maintains a pool of encoding models as {*ℋ*_1_(·), *ℋ*_2_(·),…, *ℋ*_*M*_(·)}. Given incoming neural signals *y*_*k*_, DyEnsemble weighs the models in the pool by the Bayesian likelihood to *y*_*k*_, and assembles *h*_*k*_(·)in a Bayesian averaging rule. Here, we have obtained the model pool with the TFC algorithm, with each model representing a specific state. Therefore, the DyEnsemble model can facilitate state-specific decoding by first inferring the state identity and then adaptively switching to the proper model.

Specifically, considering a time-series of neural signals *y*_0:*k*_, and the decoding problem involves estimating the kinematic state *x*_*k*_ at time step *k*. The posterior distribution of the state can be specified by:

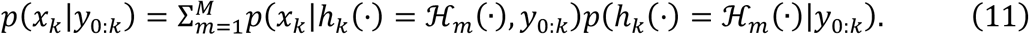

Here, *p*(*x*_*k*_|*h*_*k*_(·)= *ℋ*_*m*_(·), *y*_0:*k*_)represents the kinematic state posterior estimated by the model *ℋ*_*m*_(·)at time *k*, and *p*(*h*_*k*_(·)= *ℋ*_*m*_(·)|*y*_0:*k*_)denotes the posterior probability of selecting the encoding model *ℋ*_*m*_(·)at time *k*, which can be computed as follows:

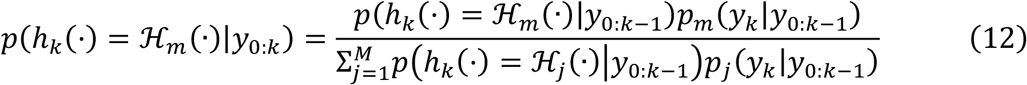

where *p*(*h*_*k*_(·)= *ℋ*_*m*_(·)|*y*_0:*k*−1_)represents the prior probability of choosing model *ℋ*_*m*_(·)at time *k*, while *p*_*m*_(*y*_*k*_|*y*_0:*k*−1_)is the marginal likelihood of choosing model *ℋ*_*m*_(·)at time *k*. The marginal likelihood represents the confidence or reliability of a particular model’s prediction.

The prior probability at time *k* can be recursively expressed as the posterior probability at time *k* − 1, with a sticking factor *α*:

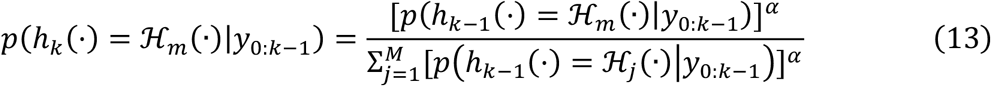

where *p*(*h*_*k*−1_(·)= *ℋ*_*m*_(·)|*y*_0:*k*−1_)is the posterior probability of choosing encoding model *ℋ*_*m*_(·)at time *k* − 1. The parameter *α* ∈ (0,1)represents the sticking factor, where a higher value leads to smoother changes in the model weights.

And the marginal likelihood can be computed as follows:

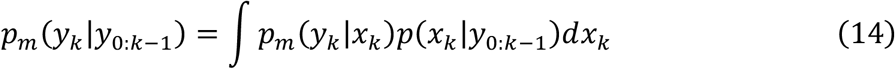

where *p*_*m*_(*y*_*k*_|*x*_*k*_)is the likelihood of model *ℋ*_*m*_(·)for a specific kinematic state *x*_*k*_. The DyEnsemble model can be solved using a particle filtering algorithm.

### Statistics

We used two statistical tests to evaluate the significance of the results. In Fig. 1i, we applied the Wilcoxon matched-pairs signed rank test (two-tailed), with n = 17 sessions. The P values are 1.526E-5 and 1.526E-5 for the RMSE and R^2^ respectively. In Fig. 2h-j, we applied the Wilcoxon matched-pairs signed rank test (two-tailed), with n = 179, n = 64, and n = 115, for 2h, 2i, and 2j, respectively. The P values for 2h, 2i, and 2j are 2.61E-54, 1.084E-19, and 4.815E-35, respectively. In Fig. 3e-f, we applied the Mann-Whitney test (two-tailed). For Fig. 3e, n = 64 and 115, with P = 1.97E-20. For Fig. 3f, n = 64 and 115, with P = 0.08134.

## Acknowledgments

We thank Xiang Li, Xufei Li, Jianfeng Yan, Hao Wu, Jianhua Qiao, and Yaoyao Hao for the technical support. This work was supported by grants from the National Key R&D Program of China (2018YFA0701400), the National Natural Science Foundation of China (62336007, 62276228), Key R&D Program of Zhejiang (2022C03011), and the Starry Night Science Fund of Zhejiang University Shanghai Institute for Advanced Study (SN-ZJU-SIAS-002).

## Author contributions

Y.Q. designed the experiment, proposed the TFC algorithm, collected and analyzed the data, and wrote the paper. X.Z. contributed to data analysis, conducted simulations, and wrote the paper. X.X. contributed to the experimental system, collected and analyzed the data, and wrote the paper. X.Y. contributed to data collection. N. D. and H.W. contributed to the manuscript writing. K.X., J.Z., and J.Z. contributed to the experiment design and implementation. Y.W. supervised the project, contributed to the experiment design and manuscript writing.

