## Extended Data Figures for "Human Motor Cortex Encodes Complex Handwriting Through a Sequence of Primitive Neural States"

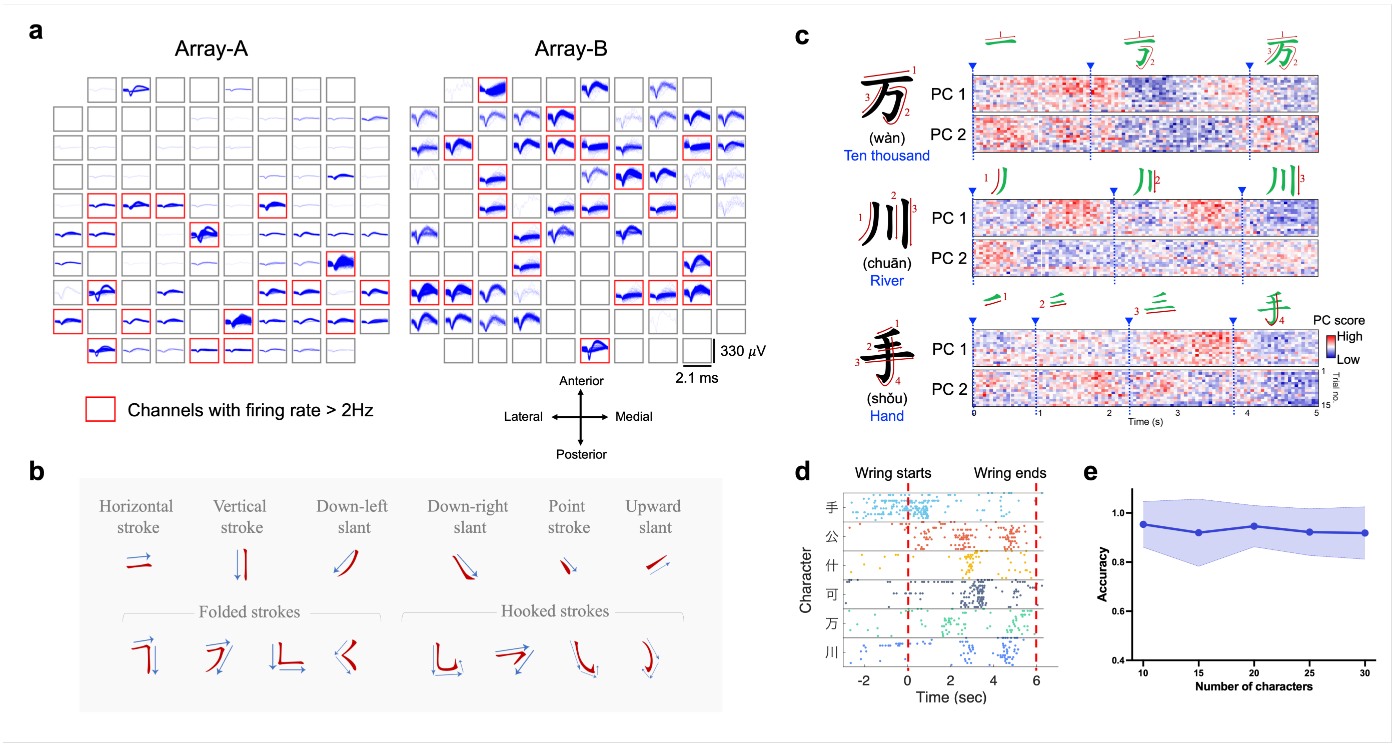


**Extended Data Fig. 1 | Example of neural signal recordings and the neural representation of handwriting. a,** Illustration of spiking activity recorded from each microelectrode array during a 3-minute window, captured on the 1402$^{nd}$ day after array implantation. **b,** Examples of strokes in Chinese writing systems. **c,** Neural activity in the top 2 principal components is shown for three example characters, 15 repetitions for each character. The color scale is normalized within each panel separately for visualization. **d,** Raster plot depicting the firing pattern of an example neuron across 15 repetitions when writing various Chinese characters. Each character begins at the 0$^{th}$ second and ends at the 6$^{th}$ second. **e,** Assessment of character classification performance using multi-channel neural signals with a linear Support Vector Machine (SVM) classifier. A total of 18 sessions, each containing 30 characters in 3 repetitions, were included. For each session, 10-30 characters were randomly selected with all repetitions, and a leave-one-trial-out test was applied. The results were averaged across all sessions, with the shaded regions showing the standard deviation.

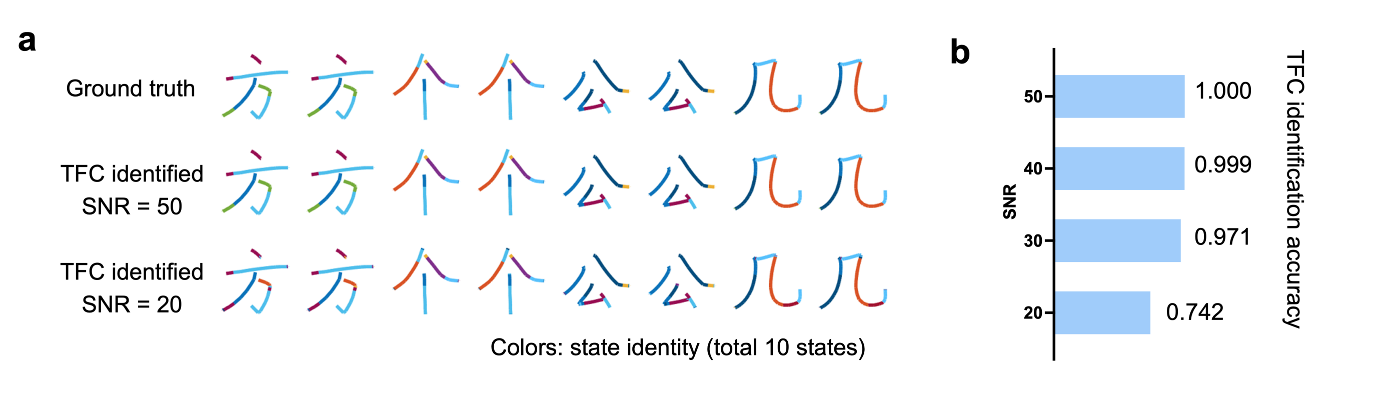


**Extended Data Fig. 2 | Evaluation of the TFC algorithm in writing state identification. a,** Simulation experiment assessing the TFC algorithm's efficacy in correctly identifying distinct mode (state) and mode (state) switch processes during handwriting (see details in Supplementary Materials). **b,** State identification accuracy under varying signal-to-noise ratios (SNR).

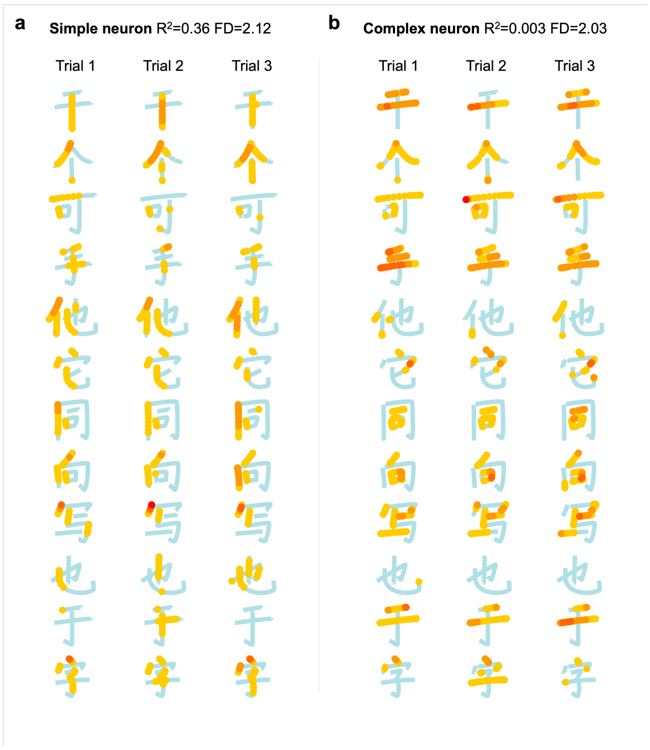


**Extended Data Fig. 3 | Examples of neuronal responses during handwriting, for both simple- and complex-tuning neurons.** **a,** Illustrative examples showcasing the neural response of a simple-tuning neuron. The neuron demonstrates a consistent preferred direction of down left, while it only selectively fires at part of down left trajectories. This neural activity can be explained by a state-dependent model, where each state shares a consistent preferred direction with diverse modulation depth. **b,** Similar to (**a**), but with a complex-tuning neuron, this neuron exhibits divergent preferred directions and fires in both leftward, rightward, and downward trajectories. This neural activity can be explained by a state-dependent directional tuning model where the preferred direction shifts across states.

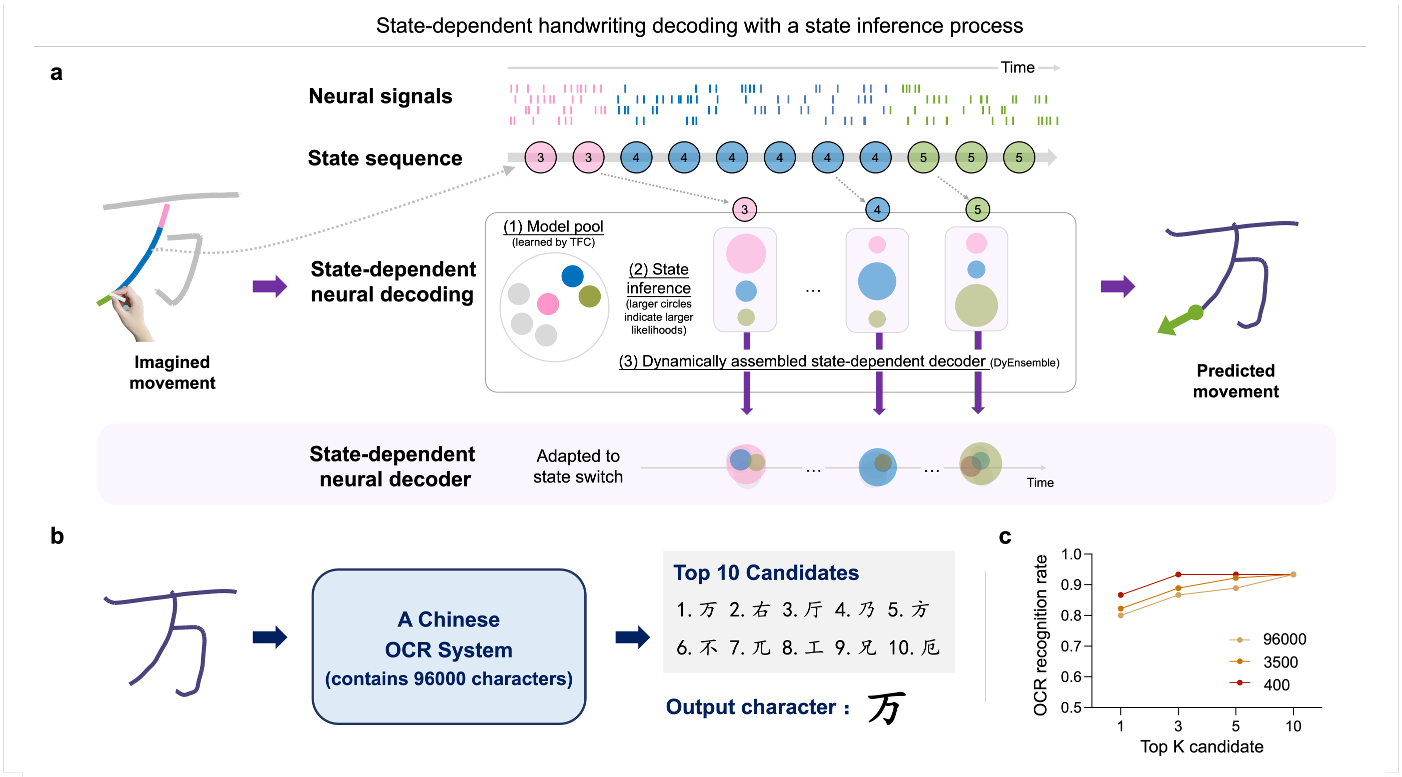


**Extended Data Fig. 4 | State-dependent decoding model improves the performance of handwriting trajectory prediction. a,** Diagrammatic representation of the state-dependent decoding process during handwriting (DyEnsemble). Utilizing encoding models for each state established through TFC, DyEnsemble dynamically infers the state and adaptively assembles a state-specific decoder in real-time based on the incoming neural signals. This approach allows for adaptive switching between decoding models along with state switches. **b.** The decoded handwriting trajectories are then fed into a commercial Chinese Optical Character Recognition (OCR) system (with a 96000 vocabulary size) to test the readability, with an exemplar session. The OCR outputs the candidates of characters for each handwriting trajectory. We evaluate how many times the correct character is within the Top K candidates as the Top K accuracy. **c,** The Top K character recognition accuracy of the OCR tool under varying vocabulary sizes.
