## Supplementary Information for "Human Motor Cortex Encodes Complex Handwriting Through a Sequence of Primitive Neural States"

### 1 Data collection sessions

We scheduled 2-3 experiment days in a week, and each experimental day contained 2-3 sessions. During a session, the participant sat in a wheelchair in an upright position, with a pillow placed to support his head and neck. His hand rested on a dinner board in front of him. A computer monitor was placed about 1 meter in front of him, indicating which character to write. The experimental sessions used for analysis in this work are listed in Table M1. The sessions used in each figure are listed in Table M2.

Table M1. Data sessions included in this study.

| Session | $N_C$ | $N_R$ | Characters |
|---|---|---|---|
| 220601-S1 | 30 | 3 | 成吃此的定对多法还好和会看来里没那你年如时是我行学永有者这作 |
| 220601-S2 | 30 | 3 | 成吃此的定对多法还好和会看来里没那你年如时是我行学永有者这作 |
| 220607-S1 | 30 | 3 | 把本但当地点动而发分过后话回活间见经开老们身世所同位用知总走 |
| 220607-S2 | 30 | 3 | 把本但当地点动而发分过后话回活间见经开老们身世所同位用知总走 |
| 220609-S1 | 30 | 3 | 爱被常到道得第都高给国果孩家理美面其起前亲使说她西现些样因种 |
| 220609-S2 | 30 | 3 | 爱被常到道得第都高给国果孩家理美面其起前亲使说她西现些样因种 |
| 220610-S1 | 30 | 3 | 不出次从大儿尔方个公几可名女去什生手他天头外为问无也在长中自 |
| 220610-S2 | 30 | 3 | 不出次从大儿尔方个公几可名女去什生手他天头外为问无也在长中自 |
| 220623-S4 | 30 | 3 | 报并场处传风告各更光合何画欢机近决军空拉利论罗妈男品求全任实 |
| 220623-S5 | 30 | 3 | 报并场处传风告各更光合何画欢机近决军空拉利论罗妈男品求全任实 |
| 220630-S1 | 30 | 3 | 变表别读房放非该花或结界金觉科苦快连林流命母呢轻师思体条往物 |
| 220630-S2 | 30 | 3 | 变表别读房放非该花或结界金觉科苦快连林流命母呢轻师思体条往物 |
| 220701-S1 | 30 | 3 | 安必边布车达代电东夫关化即加交克吗民目内平却让色声失始受书司 |
| 220701-S2 | 30 | 3 | 安必边布车达代电东夫关化即加交克吗民目内平却让色声失始受书司 |
| 220707-S1 | 30 | 3 | 巴白比产打反父火及记今乐立马片气切认水死四岁太听万王五向写由 |
| 220707-S2 | 30 | 3 | 巴白比产打反父火及记今乐立马片气切认水死四岁太听万王五向写由 |
| 220708-S1 | 30 | 3 | 完先笑信星性许言业医音应友语元员远约月运再早则怎战找至主字坐 |
| 220708-S2 | 30 | 3 | 完先笑信星性许言业医音应友语元员远约月运再早则怎战找至主字坐 |
| 221027-S1 | 6 | 15 | 川万可什公手 |
| 230327-S2 | 8 | 5 | 浙江大学脑机接口 |

| | | | |
|---|---|---|---|
| 230511-S3 | 32 | 3 | 埋坡坏坤环理玻珅权杈杯料杆种和私科秆汉叹仪扠钗汊呔忲汰钛伏吠汏状 |
| 230724-S3 | 12 | 3 | 干个可手他它同向写也于字 |

* $N_C$: The number of characters

* $N_R$: The number of repeats

Table M2. List of all data collection sessions used for the plots

| Sessions | Figures |
|---|---|
| 220601-S1 | Fig. 1i, 2a, 2g-j, 3b, 3e, f, 4a-d, Extended Data Fig. 1e |
| 220601-S2 | Fig. 1i, 2a, 2g-j, 3b, 3e, f, 4a-c, Extended Data Fig. 1e |
| 220607-S1 | Fig. 1i, 2a, 2g-j, 3b, 3e, f, 4a-c, Extended Data Fig. 1e |
| 220607-S2 | Fig. 1i, 2a, 2g-j, 3b, 3e, f, 4a-c, Extended Data Fig. 1e |
| 220609-S1 | Fig. 1i, 2a, 2g-j, 3b, 3e, f, 4a-c, Extended Data Fig. 1e |
| 220609-S2 | Fig. 1i, 2a, 2g-j, 3b, 3e, f, 4a-c, Extended Data Fig. 1e |
| 220610-S1 | Fig. 1i, 2a, 2e, 2g-j, 3a, b, 3e, f, 4a-c, Extended Data Fig. 1e |
| 220610-S2 | Fig. 1i, 2a, 2e, 2g-j, 3a, b, 3e, f, 4a-c, Extended Data Fig. 1e |
| 220623-S4 | Fig. 1i, 2a, 2g-j, 3b, 3e, f, 4a-c, Extended Data Fig. 1e |
| 220623-S5 | Fig. 2a, 2g-j, 3b, 3e, f, 4a-c, Extended Data Fig. 1e |
| 220630-S1 | Fig. 1i, 2a, 2g-j, 3b, 3e, f, 4a-c, Extended Data Fig. 1e |
| 220630-S2 | Fig. 2a, 2g-j, 3b, 3e, f, 4a-c, Extended Data Fig. 1e |
| 220701-S1 | Fig. 1i, 2a, 2e, 2g-j, 3a, b, 3e, f, 4a-c, Extended Data Fig. 1e |
| 220701-S2 | Fig. 1i, 2a, 2e, 2g-j, 3a, b, 3e, f, 4a-c, Extended Data Fig. 1e |
| 220707-S1 | Fig. 1i, 2a, 2g-j, 3b, 3e, f, 4a-c, Extended Data Fig. 1e, 4c |
| 220707-S2 | Fig. 1i, 2a, 2g-j, 3b, 3e, f, 4a-c, Extended Data Fig. 1e |
| 220708-S1 | Fig. 1i, 2a, 2g-j, 3b, 3e, f, 4a-d, Extended Data Fig. 1e |
| 220708-S2 | Fig. 1i, 2a, 2g-j, 3b, 3e, f, 4a-c, Extended Data Fig. 1e |
| 221027-S1 | Fig. 1c, Extended Data Fig. 1c, d, Extended Data Fig. 1e |
| 230327-S2 | Fig. 1h, i |
| 230511-S3 | Fig. 2a, 2g-j, 3b, 3e, f, 4a-c |
| 230724-S3 | Fig. 2a-c, 2f-j, 3b-f, 4a-c, Extended Data Fig. 3a, b |

### 2 Neural representation of handwriting

#### 2.1 The tuning behavior of neurons (Fig. 2a)

To evaluate the tuning behavior of neurons, we assessed it using two metrics: $\mathrm{R}^2$ and FD. A total of 20 experimental sessions were included (please refer to Tables M1 and M2), involving 2850 neurons.

The $\mathrm{R}^2$ of one neuron represents the proportion of its neural variance explained by the directional tuning model:

$$P(spike) = P(spike|velocity) = b_0 + b_x v_x + b_y v_y \tag{1}$$

where $v_x$ and $v_y$ are the velocity along $x$ and $y$ axis, respectively. The $\mathrm{R}^2$ was computed by:

$$\mathrm{R}^2 = 1 - \frac{\Sigma_k \left(f_k - \hat{f}_k\right)^2}{\Sigma_k \left(f_k - \bar{f}\right)^2} \tag{2}$$

where $f_k$ is the firing rate at time step $k$ of the analyzed neuron, $\hat{f}_k$ is the firing rate estimated by the directional tuning model at time step $k$, and $\bar{f}$ is the average firing rate of this neuron.

The FD (Fisher's discriminant value) is a metric indicating one neuron's discriminative ability (or reliability) of the tuning patterns by comparing the inter-character distance to the intra-character distance of neural representations, which was defined as:

$$\mathrm{FD}_C = \frac{d_{inter}^C}{d_{intra}^C} = \frac{\Sigma_{l\in C, m\notin C}(f_l - f_m)^2}{\Sigma_{l\in C, n\in C}(f_l - f_n)^2} \tag{3}$$

$$\mathrm{FD} = \frac{1}{N_C} \Sigma_C \mathrm{FD}_C \tag{4}$$

where $C$ denotes a certain target of Chinese characters and $N_C$ denotes the total number of the target characters. $l, m, n$ are trial indexes and $\Sigma_{l\in C, m\notin C}$ means finding all $(l, m)$ pairs, which satisfy trial $l$'s target is character $C$ while trial $m$'s target is not character $C$. A high FD value indicates a high discriminative ability of neuronal response, namely consistent in repetitions while discriminative with different characters.

##### 2.2 The Character-wise rasters（Fig. 2b, c, and Extended Data Fig. 3）

To plot character-wise raster, spike counts of the example neurons were first binned into 50 ms bins. Then, max-min normalization was applied to the neuronal firing rate in bins. Next, the sequence of neural responses was plotted on the corresponding trajectory of the character. Here, the handwriting trajectory is the same as the visual instruction in the video. For clearer visualization, only strong neural responses above a certain threshold (0.2) were displayed.

##### 2.3 Character classification using neural signals (Extended Data Fig. 1d)

Extended Data Fig. 1d illustrates the classification accuracy with different numbers of characters using a linear support vector machine (SVM). Neuronal spikes during writing were binned in 200 ms bins with a stride of 50 ms. Subsequently, the binned neural signals were smoothed using a moving average operation with a window size of 10. Finally, the preprocessed neural signals from all neurons were flattened into a one-dimensional vector, and fed to an SVM classifier.

Eighteen experimental sessions (including 1620 trials) were used, each of which contained 30 characters with 3 repetitions (please refer to Tables M1 and M2). For each session, we randomly selected subsets with 10, 15, 20, 25, and 30 characters and performed neural signal classification using leave-one-out cross-validation.

#### 3 State-dependent tuning during handwriting

##### 3.1 Simulation for TFC evaluation (Extended Data Fig. 2a, b)

To validate the reliability of the TFC algorithm, we conducted experiments on simulated data first as shown in Extended Data Fig. 2a, b. The simulated experiments utilized the character set from 220610-S1 consisting of 30 Chinese characters. Initially, each character was randomly divided into multiple segments, with lengths varying between 5 and 20 bins. Subsequently, we randomly generated $N_{\text{state}} = 10$ encoding models, supposing there are 10 states in the writing process. Each segment was then randomly assigned to one encoding model, and the corresponding neural signals were generated using this model and the kinematic segment. Finally, the neural signals for each character were constructed by concatenating the neural signals generated by different encoding models, with the addition of Gaussian noise at different levels. Note that we generated three sets of neural signals for each character, with the same model assignments but random noise to mimic the real experimental settings.

Specifically, we followed the method described in study[1] to generate 10 encoding models. For each encoding model, we first generated cosine tuning curves for $d_y = 100$ simulated neurons by drawing their preferred directions (PDs) randomly from a uniform distribution over $[0, 2\pi]$. Baseline firing rates were drawn from a uniform distribution over [1, 2.5], and modulation depths were drawn from a uniform distribution over [0.005, 0.025], where these parameter ranges were determined based on observations of real data. Then the encoding model can be constructed as a linear mapping matrix $B \in \mathbb{R}^{N\times(1+d_x)}$, where $d_x = 2$ is the dimension of kinematics (writing velocities along X-axis and Y-axis), 1 denotes the dimension of bias. Here, the first column of mapping matrix $B$ is the baseline firing rate of each neuron, while the second and third column of $B$ is the sine and cosine values of neuron PDs, multiplied by the corresponding modulation depth. The above process was repeated 10 times to obtain 10 encoding models. These encoding models allow us to predict the firing rates based on the kinematic inputs.

Different levels of Gaussian white noise were added to the final constructed neural signals, according to the signal-to-noise ratio (SNR):

$$\mathrm{SNR_{dB}} = 10\log_{10}\left(\frac{P_{\mathrm{signal}}}{P_{\mathrm{noise}}}\right) = 10\log_{10}\frac{\Sigma_k y_k^2}{\Sigma_k(\tilde{y}_k - y_k)^2} \tag{8}$$

where $y_k$ is the clean neural signals at time step $k$, and $\tilde{y}_k$ is the neural signals with added noise, which can be obtained directly from the function awgn() in MATLAB. In our simulation, we set up 5 noise levels of $\mathrm{SNR}_{dB} = 10, 20, 30, 40, and\ 50$, respectively.

After generating the simulation data, we then used the TFC algorithm to identify these 10 states using $M = 10$ models. The random assignment ratio $r_{\mathrm{rand}}$ was set to 0.1, while the error smooth window size $l_{\mathrm{smooth}}$ was 1. The accuracy of model assignment was reported according to:

$$acc_{\mathcal{H}_m} = \frac{\max\left(T_{\mathcal{H}_m,1}, T_{\mathcal{H}_m,2}, \dots, T_{\mathcal{H}_m,j}, \dots, T_{\mathcal{H}_m,N_{\text{state}}}\right)}{T_{\mathcal{H}_m}} \tag{9}$$

$$acc = \frac{\sum_{m=1}^{M} acc_{\mathcal{H}_m}}{M} \tag{10}$$

where $T_{\mathcal{H}_m,j}$ represents the number of overlapping indices between the data selected by model $\mathcal{H}_m$ and the data of state $j$. $T_{\mathcal{H}_m}$ denotes the length of data selected by model $\mathcal{H}_m$.

##### 3.2 Identifying primitive states with TFC (Fig. 2d, f)

We used the TFC algorithm to analyze real data and segmented the writing process into primitive states. Fig. 2f shows the examples from 230724-S3. Before applying the TFC algorithm, the spike counts were divided into 200 ms bins using a sliding window with a step of 50 ms. A moving average of 10 bins was then applied for smoothing. The neural signals corresponding to the writing phase were extracted for analysis. The TFC algorithm was configured with $M = 10$ models, randomly assignment ratio $r_{\text{rand}} = 0.1$. The error smooth window size $l_{\text{smooth}}$ was set to 5 bins.

##### 3.3 Neural encoding loss with different model numbers in TFC (Fig. 2e)

To investigate the relationship between the neural encoding capability of the TFC algorithm and the number of specified models, we varied the number of models from 1 to 100 and plotted the resulting loss curve for neural encoding. The experiment included four sessions of data, as described in Table M1 and M2. The figure plots the average encoding loss with each model number (n = 4 sessions), with the error bar denoting the standard deviation value.

As the number of models increases, the loss initially decreases at a rapid pace and then gradually slows down, exhibiting a pattern that roughly follows a power-law distribution. To further analyze this trend, we fitted the curve using the function $Y = AX^B$. The $\mathrm{R}^2$ values obtained from the curve fitting for the four sessions were 0.951, 0.945, 0.954, and 0.939, respectively. Furthermore, we applied a logarithmic transformation (base 10) to both the X and Y axes, and observed that the data could be effectively linearly fitted.

##### 3.4 The pair-wise encoding loss across states (Fig. 3a)

In Fig. 3a, we calculated the encoding loss between each pair of states, namely the models in TFC. Our analysis was based on four experimental sessions, where each session involved the collection of 30 Chinese characters with 3 repetitions (please refer to Tables M1 and M2). We employed the leave-one-repeat-out method, resulting in a total of 12 runs of the TFC algorithm. And the reported loss was the average across these runs.

Here we calculate each pair of encoding loss $\mathcal{L}(i,j)$ between state $i$ and state $j$. The configuration of TFC is the same as in Section 3.2. The computation of $\mathcal{L}(i,j)$ involved encoding the kinematic data selected by model $\mathcal{H}_i$ using model $\mathcal{H}_j$. Subsequently, we calculated the mean squared error between the encoded neural signals and the neural signals selected by model $\mathcal{H}_i$. Then the final loss was averaged over all time bins and neurons. This process can be mathematically expressed as:

$$\mathcal{L}(i,j) = \sum_{n=1}^{d_y} \sum_{k=1}^{T_{\mathcal{H}_i}} \sqrt{\left(Y_{\mathcal{H}_i} - \mathcal{H}_j\left(X_{\mathcal{H}_i}\right)\right)^2}. \tag{11}$$

Here, $Y_{\mathcal{H}_i} \in \mathbb{R}^{d_y \times T_{\mathcal{H}_i}}$ represents the neural signals selected by model $\mathcal{H}_i$, $\mathcal{H}_j(\cdot)$ denotes the encoding function of model $\mathcal{H}_j$, and $X_{\mathcal{H}_i}$ corresponds to the kinematic data selected by model $\mathcal{H}_i$.

##### 3.5 Directional tuning curves in different states (Fig. 3c, d)

To closely examine the responses from reliably tuning neurons ($R^2 \geq 0.1$ and FD≥1.4, n=179) out of 20 experimental sessions, we plotted the tuning curves of these neurons with and without considering the states. Fig. 3c, d display four example neurons from session 230724-S3, including two simple-tuning neurons and two complex-tuning neurons. The configuration of TFC is the same as in Section 3.2.

Regarding the specifics of drawing the tuning curve, the movement kinematics during writing were meticulously partitioned into eight directional intervals (0°~45°, 45°~90°, … , 315°~360°). For each interval, we computed the average firing rate of the neuron at every corresponding time point. The resulting graph illustrates the firing rates for each angle and its subsequent 45° range. Furthermore, we applied a cosine function to fit the mean firing rates and visually represented it in the graph.

For each tuning curve, we used the thickness of the curve to reflect the data size; the thicker the curve, the more data points were included for that state. Additionally, we used the color of the curve to represent the performance of the neurons; the darker the color, the higher the $R^2$ of the neuron within that state.

### 4 Neuronal representation of state occurrence

#### 4.1 The mutual information between neural activities and movement variables (Fig. 4b)

Fig. 4b evaluates the mutual information between neuronal activities with diverse movement parameters, including position (P), velocity (V), and acceleration (A), in both x and y directions. The Px in the horizontal axis indicates the position in the x direction. We plot the mean with the standard deviation as the error bar. For the shuffled condition, we randomly shuffled the neuron index at each time point, with each time point containing a neural signal from a random neuron. Then we compute the mutual information with the shuffled neural signals.

#### 4.2 The temporal firing rates during handwriting (Fig. 4d)

To plot the firing rates of a specific neuron as shown in Fig. 4d, the spike counts were first binned into 50 ms bins without overlap. Next, the binned data was smoothed using a 5-step moving average filter. The data used for plotting the figure was from the go cue to the end of the writing. The solid line in the plot represents the trial-averaged firing rates of one Chinese character (n=3 trials). This was obtained by averaging the firing rates across multiple trials for the same character. To visualize the variability in the firing rates across trials, the shaded regions in the plot indicate the standard deviation of the mean firing rates.

### 5 State-dependent handwriting decoding

#### 5.1 Visualization of decoded writing trajectories (Fig. 1h)

We selected the four Chinese characters “脑机接口” (meaning: Brain-computer interface) to illustrate the decoding performance. Two decoders were compared, with one state-dependent decoder (DyEnsemble, see Methods for details) and a decoder without considering states (Kalman filter). Initially, the spike counts were grouped into 200 ms bins using a sliding window with a step of 50 ms. To smooth the signals, a moving average with a window size of 10 bins was applied. For decoding, we employed a cross-validation approach known as leave-one-repeat-out. This means that the preprocessed data was divided into training sets and test sets according to the number of times each character was repeated. In each iteration, we used one repeat for test and the remaining repeats for training.

For the DyEnsemble decoder, we first utilized the TFC algorithm to identify $M = 10$ states and their associated models in the training data. The parameter $l_{\mathrm{smooth}}$ was set to 10, and the early-stopping threshold $\beta$ was set to 0.001. Subsequently, we used the dynamic ensemble algorithm to calculate the weights for each model and predict the kinematic states based on the neural signals at each time step during testing. The particles used in the algorithm were unique kinematic states from the training set, and the sticking factor $\alpha$ was set to 0.1.

For the Kalman filter (KF), we employed a velocity Kalman filter with a linear-Gaussian state-space model[2]:

$$x_k = Ax_{k-1} + b + \varsigma_{k-1}, \tag{12}$$

$$y_k = Hx_k + \varepsilon_k \tag{13}$$

where $k$ denotes the time step index, $x_k \in \mathbb{R}^{d_x}$ is the kinematic state and $d_x = 2$ is the dimensions of $x_k$, including the writing velocities in X-axis and Y-axis. $y_k \in \mathbb{R}^{d_y}$ is the neural measurement and $d_y$ is the dimensions of $y_k$, determined by the number of neurons recorded in this session. $\varsigma_k \sim N(0, \sigma_\varsigma^2)$, $\varepsilon_k \sim N(0, \sigma_\varepsilon^2)$ are state transition noise and measurement noise. For each stroke, we adopted the standard stroke start position. The stroke trajectory was obtained by integrating the decoded velocities.

#### 5.2 Evaluation metrics of handwriting decoding performance (Fig. 1i)

To generate the statistical results for Fig. 1i, the decoding performance of the Kalman filter and the DyEnsemble decoder was evaluated using 17 experimental sessions, consisting of 1480 trials with 488 targets.

Three evaluation metrics were used for the decoded velocity: root mean squared error (RMSE), and coefficient of determination ($R^2$). The formulas for these metrics are as follows:

$$\text{RMSE} = \sqrt{\frac{1}{T}\Sigma_{k=1}^{T}(x_k - \hat{x}_k)^2} \tag{14}$$

$$R^2 = 1 - \frac{\Sigma_k (x_k - \hat{x}_k)^2}{\Sigma_k (x_k - \bar{x})^2} \tag{16}$$

In these formulas, $x_k$ represents the velocity at time step $k \in [1, T]$, $\hat{x}_k$ represents the velocity estimated by the decoder at time step $k$, and $\bar{x}$ is the average velocity of all time steps.

To calculate the final performance of each session, the metrics were averaged over all replicates. In the graph, each data point represents the decoding result of a session (n=17). To assess the significance of the decoding performance between the two algorithms, a Wilcoxon matched-pairs signed rank test was performed, with $P < 10^{-4}$ in terms of RMSE and $R^2$.

#### 5.3 Character recognition accuracy with OCR (Extended Data Fig. 4c)

To evaluate the legibility of the decoding trajectory, we utilized a commercial Optical Character Recognition (OCR) tool[3]. In each trial, the decoded velocities were integrated to form trajectories. These trajectories were then centered and mapped between 0 and 255 before being sent to the OCR tool. Subsequently, a list of candidate characters was obtained from the OCR tool.

To accommodate different usage scenarios, we considered three different character vocabulary sizes. These included a vocabulary of 96,000 Chinese characters, covering almost all Chinese characters, a vocabulary of 3,500 commonly used Chinese characters, and a vocabulary of 400 Chinese characters collected during our experiments. These vocabularies were used to filter the candidate character lists generated by the OCR tool, resulting in a list of refined candidate characters.

In our analysis, we computed the recognition rates for various levels of fault tolerance, namely top1, top3, top5, and top10 accuracy. To clarify, a trial is considered recognized correctly if the true target character is among the top k candidates in the refined candidate character list. The recognition rate is then calculated as the ratio of correctly recognized trials to the total number of trials.
